# Vaccination-induced rapid protection against bacterial pneumonia via training alveolar macrophage in mice

**DOI:** 10.1101/2021.05.10.443370

**Authors:** Hao Gu, Xi Zeng, Liusheng Peng, Chuanying Xiang, Yangyang Zhou, Xiaomin Zhang, Jixin Zhang, Ning Wang, Gang Guo, Yan Li, Kaiyun Liu, Jiang Gu, Hao Zeng, Yuan Zhuang, Haibo Li, Jinyong Zhang, Weijun Zhang, Quanming Zou, Yun Shi

**Author notes:** Corresponding authors, Yun Shi, West China Biopharm Research Institute, West China Hospital, Sichuan University, Chengdu, Sichuan 610041, China; (Y.S.). Quanming Zou, National Engineering Research Center of Immunological Products, Third Military, Medical University, Chongqing 400038, China; (Q. Z.). These authors contributed equally to this work (Hao Gu, Xi Zeng).

## Abstract

Vaccination strategies for rapid protection against multidrug-resistant bacterial infection are very important, especially for hospitalized patients who have high risk of exposure to these bacteria. However, few such vaccination strategies exist due to a shortage of knowledge supporting their rapid effect. Here we demonstrated a single intranasal immunization of inactivated whole cell (IWC) of *Acinetobacter baumannii* elicits rapid protection against *A. baumannii*-infected pneumonia via training of innate immune response in *Rag1*^-/-^ mice. Immunization-trained alveolar macrophages (AMs) showed enhanced TNF-α production upon restimulation. Adoptive transfer of immunization-trained AMs into naive mice mediated rapid protection against infection. Elevated TLR4 expression on vaccination-trained AMs contributed to rapid protection. Moreover, immunization-induced rapid protection was also seen in *Pseudomonas aeruginosa* and *Klebsiella pneumoniae* pneumonia models, but not in *Staphylococcus aureus* and *Streptococcus pneumoniae* model. Our data reveal that a single intranasal immunization induces rapid and efficient protection against certain Gram-negative bacterial pneumonia via training AMs response, which highlights the importance and the possibility of harnessing trained immunity of AMs to design rapid-effecting vaccine.

## Introduction

The multidrug-resistant (MDR) bacteria, including *Acinetobacter baumanni, Pseudomonas aeruginosa, Klebsiella pneumoniae, Escherichia coli*, and *Staphylococcus aureus*, pose a great threat to global public health (Tacconelli, 2017). Pneumonia caused by MDR bacteria is a major cause of morbidity and mortality, especially in hospitalized patients (Gonzalez-Villoria & Valverde-Garduno, 2016; Micek et al., 2015; Zilberberg, Nathanson, Sulham, Fan, & Shorr, 2016). The continuing spread of antimicrobial resistance has made treating MDR bacterial pneumonia extremely difficult. Vaccination has been proposed as a promising strategy for controlling MDR bacterial infections (Jansen, Knirsch, & Anderson, 2018; Rappuoli, Bloom, & Black, 2017; Williams, 2007). Current vaccination strategies usually require multiple injections weeks or months apart, which limit them to rapidly prevent infections for inpatients. However, hospitalized patients have an especially high risk for exposure to MDR bacteria (Pachon & McConnell, 2014). Therefore, rapid efficacy induced by vaccination is vital for vaccine development against MDR bacteria (Pachon & McConnell, 2014).

Induction of immunological memory is the central goal of vaccination. Immunological memory protects against infections by enabling a quicker and stronger immune response to a previously encountered antigen (Farber, Netea, Radbruch, Rajewsky, & Zinkernagel, 2016). Classically, immune memory is thought to be exclusively mediated by adaptive T and B cell responses. These responses are highly specific to antigen, but take days or weeks to become effective. Another part of the immune system, innate immune response, provides an initial, relatively nonspecific response to infection within hours to days without immunological memory. However, in the past decade, evidence has emerged showing that innate immune cells such as monocytes, macrophage, and NK cells can also build long-term memory through epigenetic and metabolic reprogramming of cells. This memory termed “trained immunity” or “trained innate immunity,’’ produces hyperresponsiveness upon re-stimulation in these cells (Netea et al., 2016; Netea & Joosten, 2018; Netea, Quintin, & van der Meer, 2011). The rapidity of innate immune response leads us to speculate that trained innate immunity might effectively serve as the underlying mechanism for vaccination-induced rapid protection.

Here, we demonstrated that a single intranasal immunization of inactivated whole cell (IWC) induced rapid and efficient protection against certain Gram-negative bacterial pneumonia, which was dependent on trained innate immunity mediated by alveolar macrophages (AMs). These findings highlight the possibility to harness the trained immunity of AMs to design a vaccine with rapid efficacy against pulmonary infection.

## Results

### Rapid protection against *A. baumannnii* pneumonia by a single intranasal vaccination

Mice were immunized intranasally (i.n.) with an inactivated whole cell (IWC) of *A. baumannii* and infected intratracheally (i.t.) with a lethal dose of *A. baumannii* 2 days or 7 days later (Figure 1A). The control mice succumbed to the infection, whereas all IWC-vaccinated mice survived when mice were challenged 2 days or 7 days post immunization (Figure 1B, *\*\* P* < 0.01 compared to control, log-rank test). Consistent with the survival rate, bacterial burdens in lungs and blood of vaccinated mice were significantly lower than those in the control group at 24 hours post infection (hpi) (Figure 1C, ** *P* < 0.01, *** *P* < 0.001, ordinary one-way ANOVA). Histopathology of lung tissues showed reduced lung damage and decreased inflammatory cells infiltration in IWC-immunized mice (Figure 1D). When challenging the mice at day 7 post immunization, the pro-inflammatory cytokines of IL-6 in lungs and serum levels of IL-6 and TNF-α (Figure 1E and Figure 1-figure supplement 1, **** *P* < 0.0001, unpaired *t* test.) in IWC-vaccinated group were significantly lower than those in the control group at 24 hpi. Expression of chemokines *Cxcl1, Cxcl2, Cxcl5, Cxcl10*, and *Ccl2* were also significantly reduced in the lungs of vaccinated mice at 24 hpi (*Figure 1F*, * *P* < 0.05, ** *P* < 0.01, and **** *P* < 0.0001, unpaired *t* test.). Inflammatory cells in the lungs were detected by the flow cytometry and gating strategy was shown in *Figure 1-figure supplement 1B*. The results showed that the number of neutrophils in lungs of vaccinated mice was significantly lower than that of control group at 24 hpi (Figure 1G and Figure 1-figure supplement 1C. *\*\* P* < 0.01, unpaired *t* test). Collectively, these findings indicate a single intranasal immunization with IWC elicits rapid and complete protection against pulmonary *A. baumanniii* infection.

**Figure 1.**
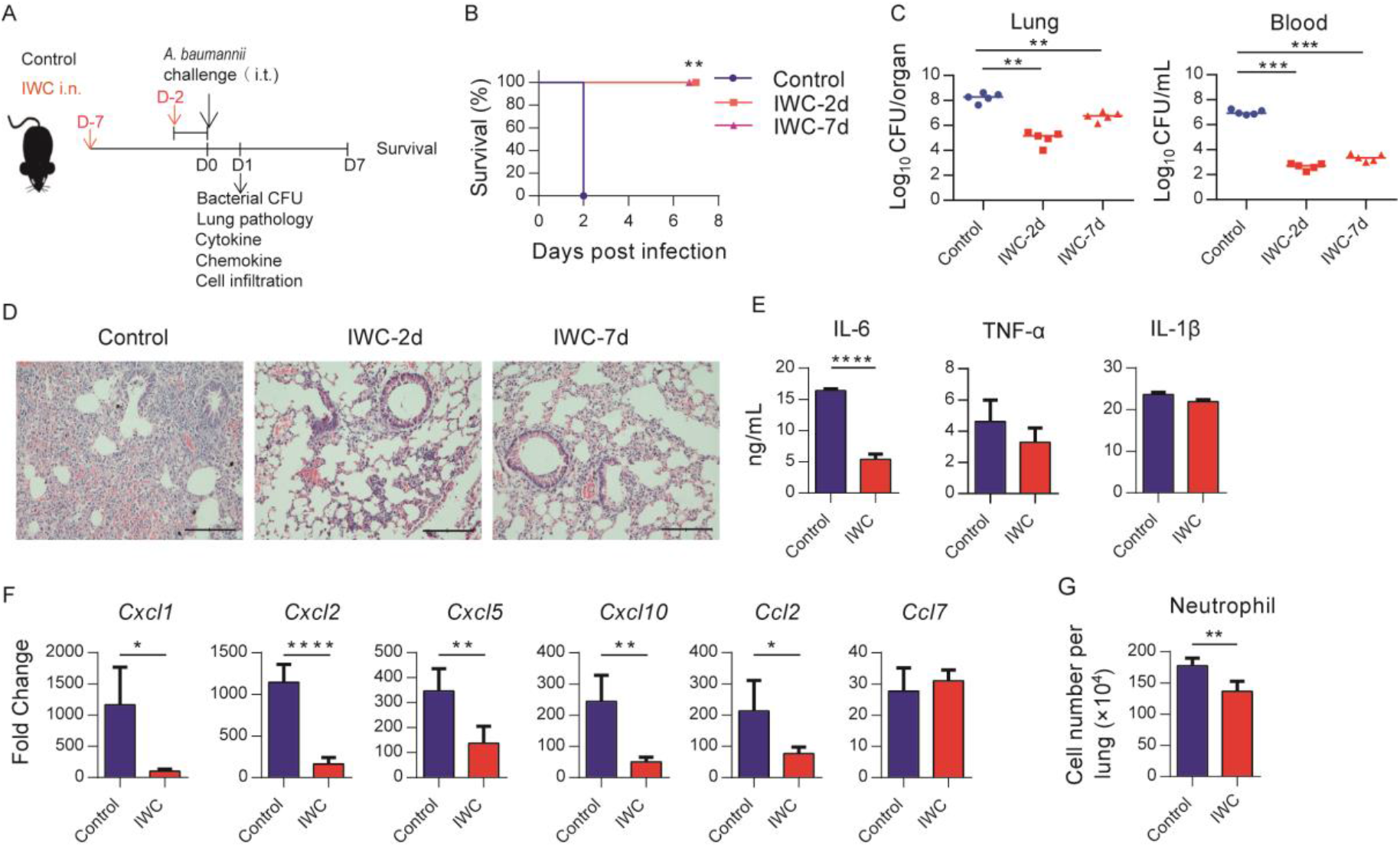
Rapid protection against *A. baumannnii* pneumonia by a single intranasal vaccination. **(A)** Schematic diagram of the experimental procedure. C57BL/6 mice were immunized intranasally (i.n.) with inactivated whole cell (IWC) of *A. baumannii* and challenged intratracheally (i.t.) with *A. baumannii* at day 2 (IWC-2d) or day7 (IWC-7d) after immunization (n=5/group). (**B**) Survival of mice was recorded for 7 days. \*\**P* < 0.01 determined by log-rank test. (**C**) Bacterial burdens in lungs and blood at 24 hour post infection (hpi) was determined. Each plot represents one mouse. The line indicates the median of the data. \*\**P* < 0.01, \*\*\**P* < 0.001 evaluated by ordinary one-way ANOVA followed by Tukey’s multiple comparisons test. (**D**) Representative histopathologic images of lungs at 24 hpi. Scale bars: 100 μm. (**E-G**) IWC-immunized mice were challenged at day 7 and were sacrificed at 24 hpi. (E) Levels of inflammatory cytokines in the lungs were detected by ELISA. (**F**) Transcriptional levels of chemokines in the lungs were detected by real-time PCR. (**G**) Numbers of neutrophils in the lungs were detected by flow cytometry. Data are mean ± SD. n=4-5 mice/group. For (**E**) to (**G**), * *P* < 0.05, ** *P* < 0.01, and **** *P* < 0.0001, determined by two-tail unpaired *t* test. Data are representative of at least two independent experiments.

### Rapid immune memory induced by a single intranasal vaccination

Immunological memory is defined as functionally enhanced, quicker, and more effective response to pathogens that have been encountered previously. This is the basis of successful vaccines against subsequent infections. To assess whether the IWC-induced rapid protection is a result of immunological memory or a result of the persistent activation of innate immune responses, we measured the dynamic immune response after immunization of IWC from day 0 to day 7. In response to intranasal immunization of IWC, levels of TNF-α and IL-6 in lungs increased from day 1 to day 4 and completely declined to baseline by day 5 (Figure 2A), indicating that the host response rapidly primed and rest 5 days later. When we challenged the mice with *A. bauammnii* on day 7 after vaccination and assessed the cytokine levels in lungs early at 2 hpi, we found that TNF-α but not IL-6 and IL-1β levels was significantly higher from IWC-immunized mice than those from control mice (Figure 2B, ***, *P* < 0.001, ordinary two-way ANOVA). Further, mRNA levels of *Cxcl1, Cxcl2, Cxcl5, Cxcl10*, and *Ccl2* were higher in vaccinated mice than in control mice at early 2 hpi (Figure 2C, \**P* < 0.05; \*\**P* < 0.01, \*\*\*\**P* < 0.0001, ordinary two-way ANOVA). Meanwhile, consistent with the increased chemokine expression, vaccinated mice had significantly higher numbers of neutrophils and monocytes in their lungs than did control mice at 4 hpi (Figure 2D, \*\**P* < 0.01, \*\*\*\**P* < 0.0001, ordinary two-way ANOVA). These results indicate IWC immunization induces quicker, enhanced responses upon *A. baumannii* challenge on day 7 after vaccination. Vaccination-induced protection against *A. baumannii* on day 7 after vaccination is an enhanced recall response of immune response.

**Figure 2.**
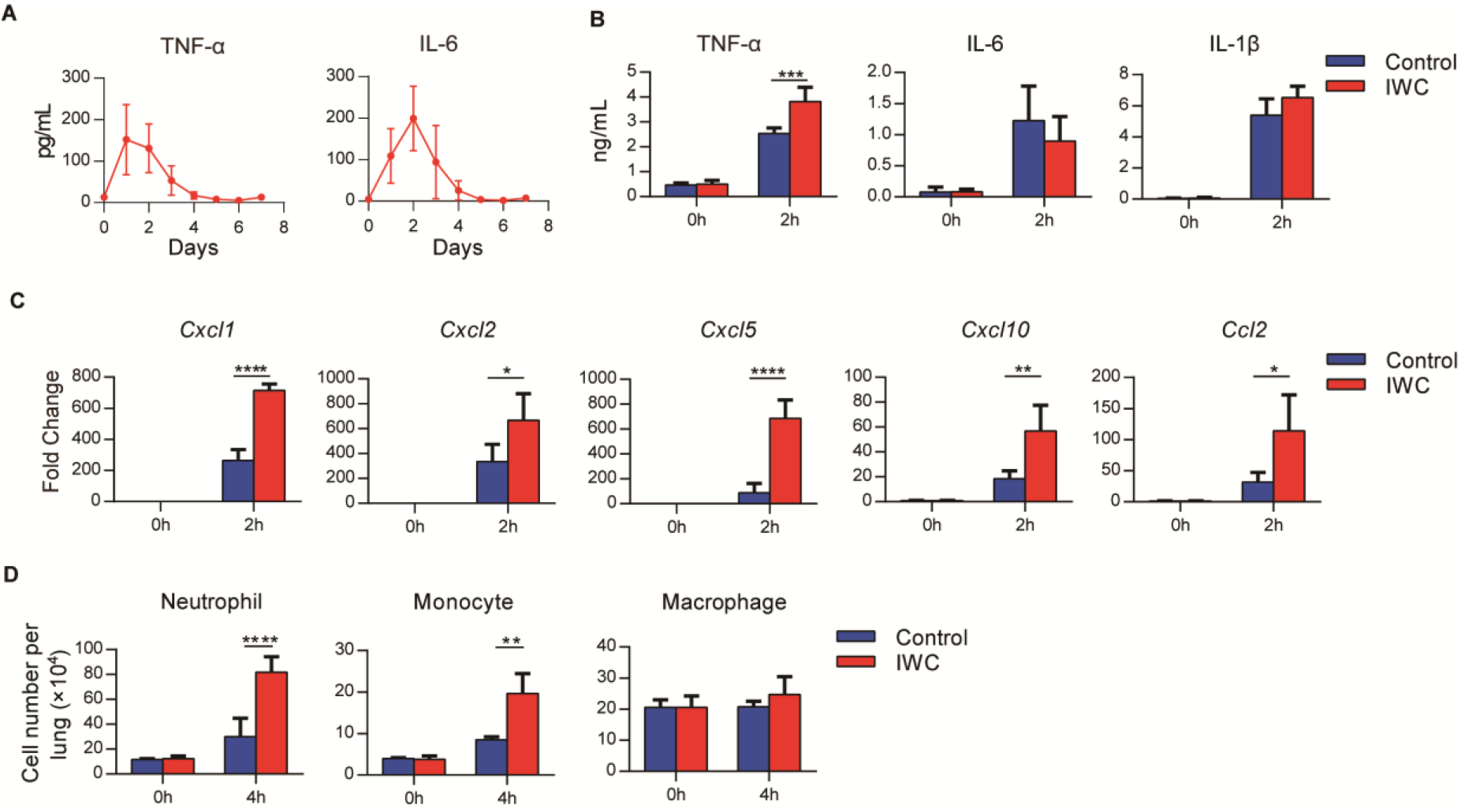
Rapid immune memory induced by a single intranasal vaccination. (**A**) Dynamic responses of TNF-α and IL-6 in the lungs of *A. baumannii* IWC-immunized mice (n = 3 per timepoint). (**B-D**) IWC-immunized mice were challenged i.t. with *A. baumannii* at day 7 after immunization. (**B**) Levels of TNF-α, IL-6, and IL-1β at 0 h and 2 hpi in the lungs were measured by ELISA. (**C**) Transcriptional levels of chemokines in the lungs at 0 h and 2 hpi were assessed by real-time PCR. (D) Numbers of neutrophils, monocytes, and macrophages in the lungs of mice were determined by flow cytometry. Data are presented as mean ± SD. (n=3-4 mice/group). \**P* < 0.05; \*\**P* < 0.01, \*\*\**P* < 0.001, \*\*\*\**P* < 0.0001, ordinary two-way ANOVA. Data are representative of two independent experiments.

### Vaccination-induced rapid protection is dependent on trained innate immunity

To further explore the potential mechanism of rapid effect of vaccination, *Rag1*^-/-^ mice (which lack mature T and B cells) were immunized i.n. with IWC and then challenged with a lethal dose of *A. baumannii* to determine which part of immune response is responsible for rapid protection. Results showed IWC provided rapid and effective protection both in WT and *Rag1*^-/-^ mice at day 7 after immunization (Figure 3A, ** *P* < 0.01 compared to control, log-rank test. There is no significant difference between WT and *Rag1*^*-/-*^ mice in terms of survival of the IWC-immunized group (Figure 3A, P=0.30, log-rank test). RNA sequencing analyses (RNA-seq) of lung tissue at day 7 after immunization and 24 h after *A. baumannii* challenge revealed significant differentially expressed genes (DEGs) between transcriptional profiles of vaccinated *Rag1*^-/-^ mice and those of control *Rag1*^-/-^ mice (Figure 3B and Figure 3-figure supplement 1A). There were a total of 2401 DEGs between these two groups; 1084 upregulated genes and 1317 downregulated in the vaccinated mice (Figure 3-figure supplement 1B). Gene ontology (GO) analysis of DEGs revealed that genes associated with inflammatory response, response to molecule of bacterial origin, and response to liposaccharide were significantly downregulated in vaccinated mice at 24 hpi (Figure 3C and Figure 3-figure supplement 1C). The expression of inflammation related genes including *Il6, Cxcl1, Cxcl2, Cxcl10*, and *Ccl2* were notably lower in vaccinated-*Rag1*^-/-^ mice than those in control mice at 24 hpi (Figure 3D and E and Figure 3-figure supplement 1D, \*\**P* < 0.01, unpaired *t* test). These data indicate that IWC immunization induces rapid protection in *Rag1*^-/-^ mice, highlighting the role of innate immune response in vaccination-induced rapid protection. We also analyzed the dynamic transcriptional response to IWC immunization in *Rag1*^-/-^ mice and found that the innate immune response was activated at day 2 and rested at day 7 (Figure 3F), which has the similar pattern to that in control mice (Figure 2). Upon *A. baumannii* challenge at day 7 after immunization, the immunized mice exhibited different responses to those unimmunized mice (Figure 3-figure supplement 1E). These results indicate IWC immunization induces a trained feature of innate immune response, which is critical for vaccination-induced rapid protection.

**Figure 3.**
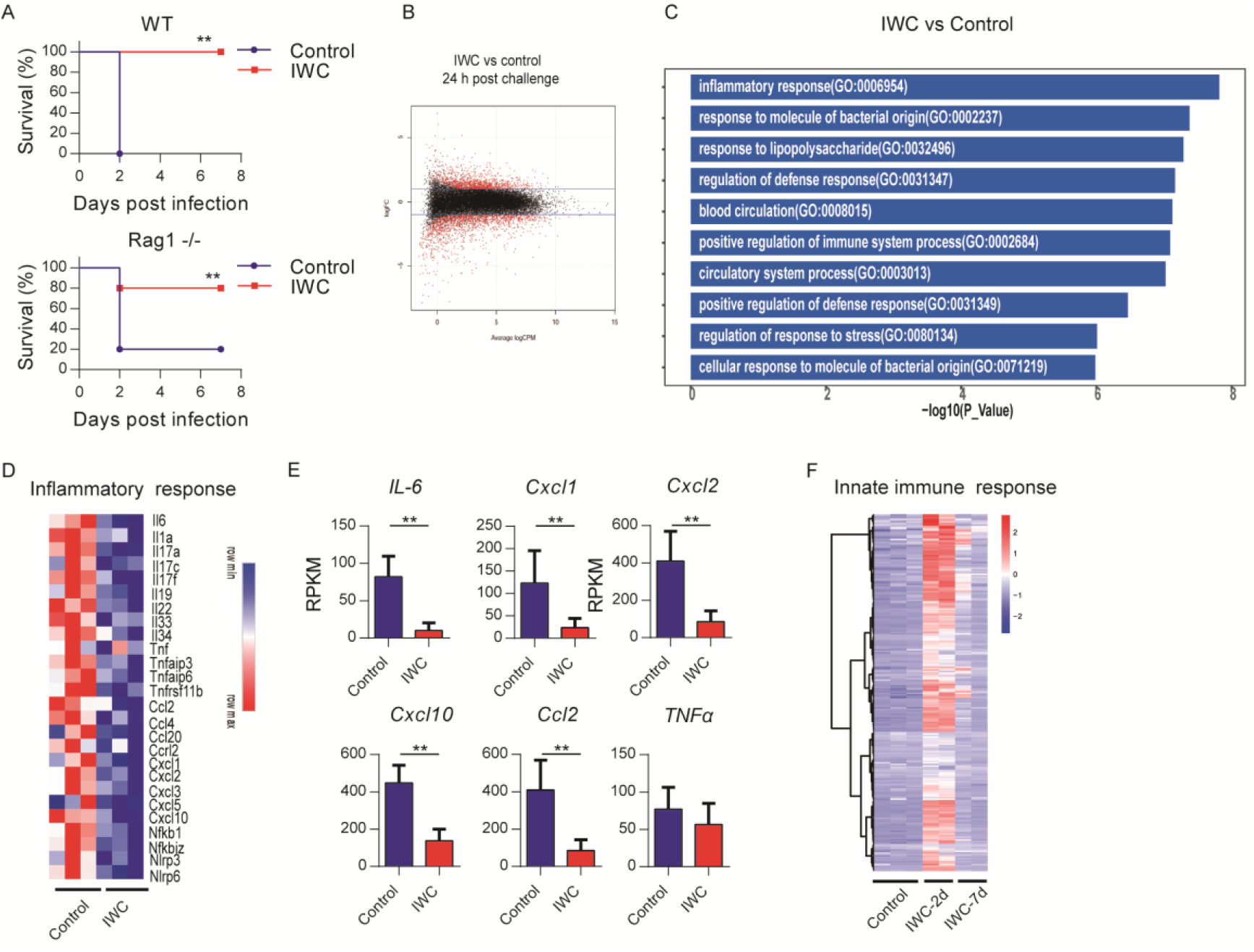
Trained innate immunity mediates vaccination-induced rapid protection. (**A**) Survival of WT and *Rag1*^-/-^ mice immunized i.n. with IWC or PBS, challenged i.t by lethal *A. baumannii* 7 days later (n=5/group for WT mice, n=10 /group for *Rag1*^-/-^ mice). ** *P* < 0.01 compared to control calculated by log-rank test. Data are representative of two independent experiments. (B) MA plot of the DEGs of IWC-immunized mice vs control mice at 24 hpi. X-axis represents average counts-per-million (logCPM) and Y-axis represents log fold-changes (logFC) in IWC-immunized mice vs control mice. The blue line is the threshold, logFC > 1 means upretulation and logFC < −1 means downregulation. (**C**) Top 10 GO enrichment terms of downregulated DEGs in the IWC-immunized group at 24 hpi. (**D**) Heatmap of DEGs related to inflammatory response was shown. False discovery rate (FDR) < 0.05. (**E**) Reads per kilobase per million reads (RPKM) of inflammatory and chemokine genes. Data are mean ± SD. \*\**P* < 0.01 determined by two-tailed unpaired *t* test. (**F**) The heatmap of innate immune response related genes (GO_0045087) of lung samples from control, IWC-immunized at day 2 (IWC-2d), and day 7 (IWC-7d) in *Rag1*^-/-^ mice.

### Trained immunity of alveolar macrophages mediates vaccination-induced rapid protection

Further, we analyzed the transcriptome change induced by vaccination at day 7 to identify DEGs associated with trained innate immunity. RNA-seq data showed a total of 308 DEGs in lungs of IWC-vaccinated *Rag1*^-/-^ mice at day 7 (Figure 4A and Figure 4-figure supplement 1A). The upregulated 253 DEGs were enriched to myeloid leukocyte activation (Figure 4B) and these genes were enriched to macrophage-associated genes (Figure 4-figure supplement 1B). So, we reasoned that alveolar macrophages (AMs), the predominant patrol myeloid cells in airways, might play a key role in vaccination-induced rapid protection. To test this hypothesis, we established a trained immunity model of AMs by stimulation with IWC *in vitro*. IWC-trained AMs induced an enhanced TNF-α production upon restimulation with IWC 7 days later. This result indicates that IWC trains AMs directly (Figure 4C, *P* < 0.01, unpaired *t* test.). AMs from IWC-immunized or control mice at day 7 were sorted from bronchoalveolar lavage fluid (BALF) using CD11c^+^ microbeads. Flow cytometry analysis with anti-CD11c and anti-F4/80 confirmed the purity of AMs was greater than 95% (Figure 4-figure supplement 1C). Sorted AMs were stimulated with IWC *ex vivo* for 2 h. TNF-α production in vaccinated AMs after restimulation was significantly higher than that in control AMs (Figure 4D, \**P* < 0.05, unpaired *t* test.). Thus, AMs could be trained by IWC with functional reprogramming, showing increased TNF-α production to a previously encountered stimulus. Further, the purified AMs from *A. baumannii* IWC-immunized or control mice at day 7 were adoptively transferred into the airway of naïve mice by direct intra-tracheal instillation. Upon *A. baumannii* challenge, the lungs of mice that had received transfer of IWC-primed AMs had significantly lower bacterial burdens with alleviated clinical scores (Figure 4E, *, *P* < 0.05, **,*P* < 0.01, Mann-Whitney U test)). Further treatment of IWC-immunized mice with anti-TNF-α antibody before *A. baumannii* challenge resulted in reduced protection (Figure 4F, *, *P* < 0.05, log-rank test.). These results indicate that vaccination-trained AMs mediate rapid protection against infection via enhanced TNF-α production.

**Figure 4.**
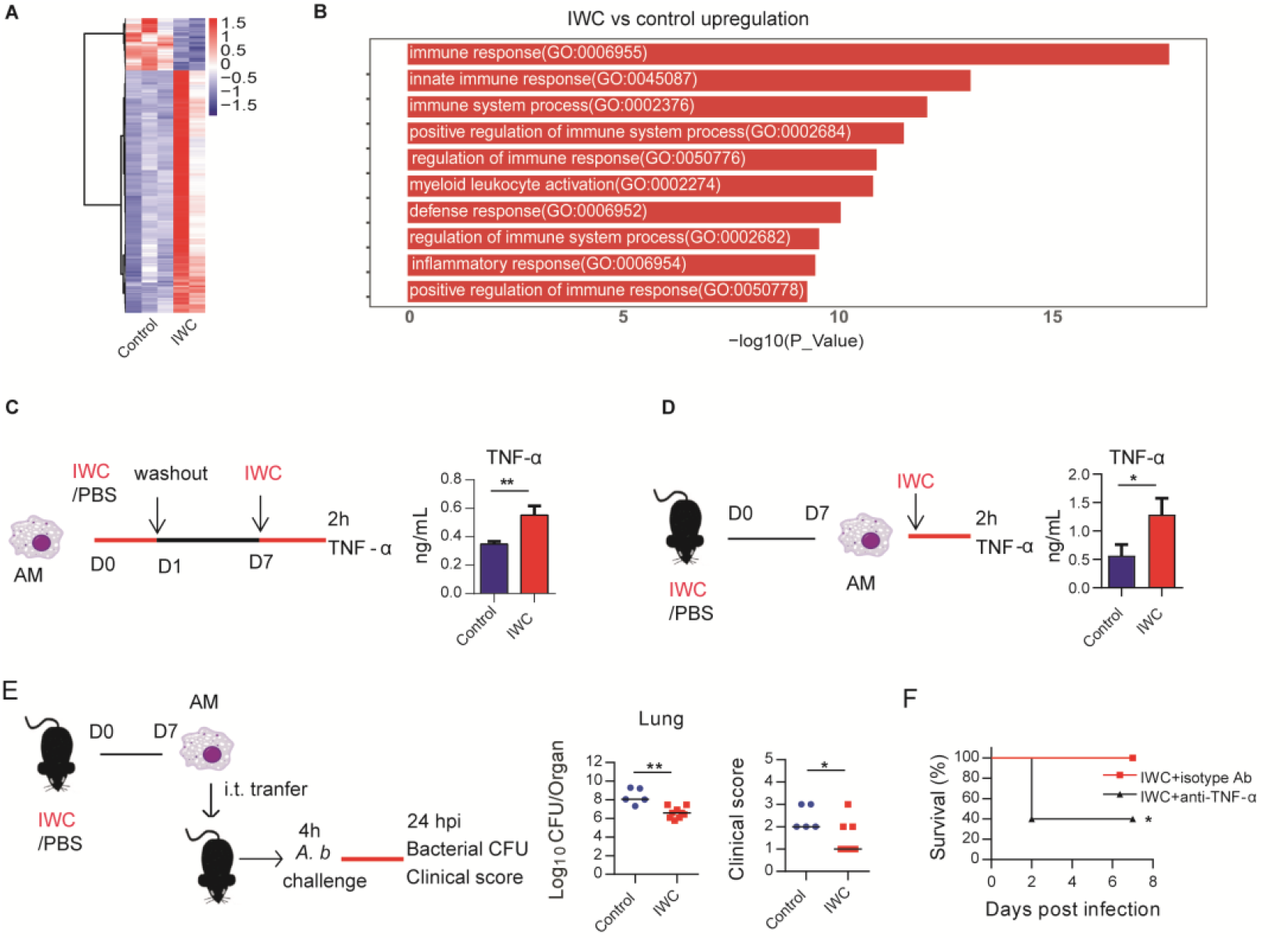
Trained immunity of AMs mediates rapid protection induced by vaccination. (**A**) Heatmap of DEGs in lungs of IWC-immunized and control *Rag1*^-/-^ mice at day 7 after immunization. (B) Top 10 GO terms of upregulated DEGs in IWC-immunized group at day 7. (**C**) *In vitro* model of IWC-trained AMs. (**D**) C57BL/6 mice were immunized with IWC and recall responses of trained AMs to IWC were evaluated *ex vivo* by detection of TNF-α production at 2 h after stimulation. For C and D, Data are mean ± SD. n=3. \**P* < 0.05, \*\**P* < 0.01, two-tail unpaired *t* test. (**E**) Schema of evaluating roles of AMs in BALF of PBS or IWC-immunized C57BL/6 mice. Bacterial burdens and clinical scores at 24 hpi were measured (n=5-9). The line represents the median. * *P* < 0.05, \*\**P* < 0.01 determined by Mann-Whitney U test. (**F**) WT mice were immunized with IWC for 7 days. Mice were treated intraperitoneally with anti-TNF-α antibody or isotype control then were challenge with lethal *A. baumannii* 1 hour later. The survival of mice was monitored (n = 5). *, *P* < 0.05, calculated by log-rank test. Data are representative of two independent experiments.

### Contribution of higher TLR4 expression on trained AMs to rapid protection

RNA-seq reveals that genes related to myeloid leukocyte activation including TLRs were significantly upregulated on day 7 after *A. baumannii* IWC immunization in *Rag1*^-/-^ mice (Figure 5A and Figure 5-figure supplement 1). These results suggest that surface molecules associated with cell activation might be markers for trained AMs. Since TLR4 plays an important role in host recognition of Gram-negative bacteria, we hypothesized that IWC-trained AMs with elevated TLR4 expression might be more sensitive for second recall activation and enhanced function. RAN-seq data showed that lung TLR4 transcript significantly increased in response to IWC immunization at day 2 and day 7 (Figure 5B, \**P* < 0.05, one-way ANOVA). The elevated TLR4 expression on BALF AMs at day 2 or day 7 after IWC immunization was also confirmed by flow cytometry (Figure 5C, \*\**P* < 0.01, one-way ANOVA). We also found TLR4 expression on AMs was elevated at day 7 after IWC training *in vitro* (Figure 5D, \**P* < 0.05, unpaired *t* test). Further, we found that the rapid protective effect of IWC-vaccination was significantly reduced in *Tlr4*^-/-^ mice than in WT mice (Figure 5E. * *P*<0.05, log-rank test). Accordingly, IWC-vaccination could not reduce bacterial burdens in lungs and blood in *Tlr4*^-/-^ mice upon *A. baumannii* challenge (Figure 5F, \**P* < 0.05, ordinary two-way ANOVA). IWC-immunization induced rapid TNF-α expression at 2 hpi and neutrophil infiltration at 4 hour post *A. baumannii* challenge were dismissed in *Tlr4*^-/-^ mice (Figure 5G and 5H, **, *P* < 0.01, ****, *P* < 0.0001, ordinary two-way ANOVA). In addition, we found that IWC-priming Ams from *Tlr4*^-/-^ mice significantly lost TNF-α secretion in response to IWC restimulation *ex vivo* (Figure 5I). These results suggest TLR4 signaling is vital for IWC-trained AMs. Taken together, these results suggest that up-regulation of TLR4 expression on trained AMs plays an important role in vaccination-induced rapid protection.

**Figure 5.**
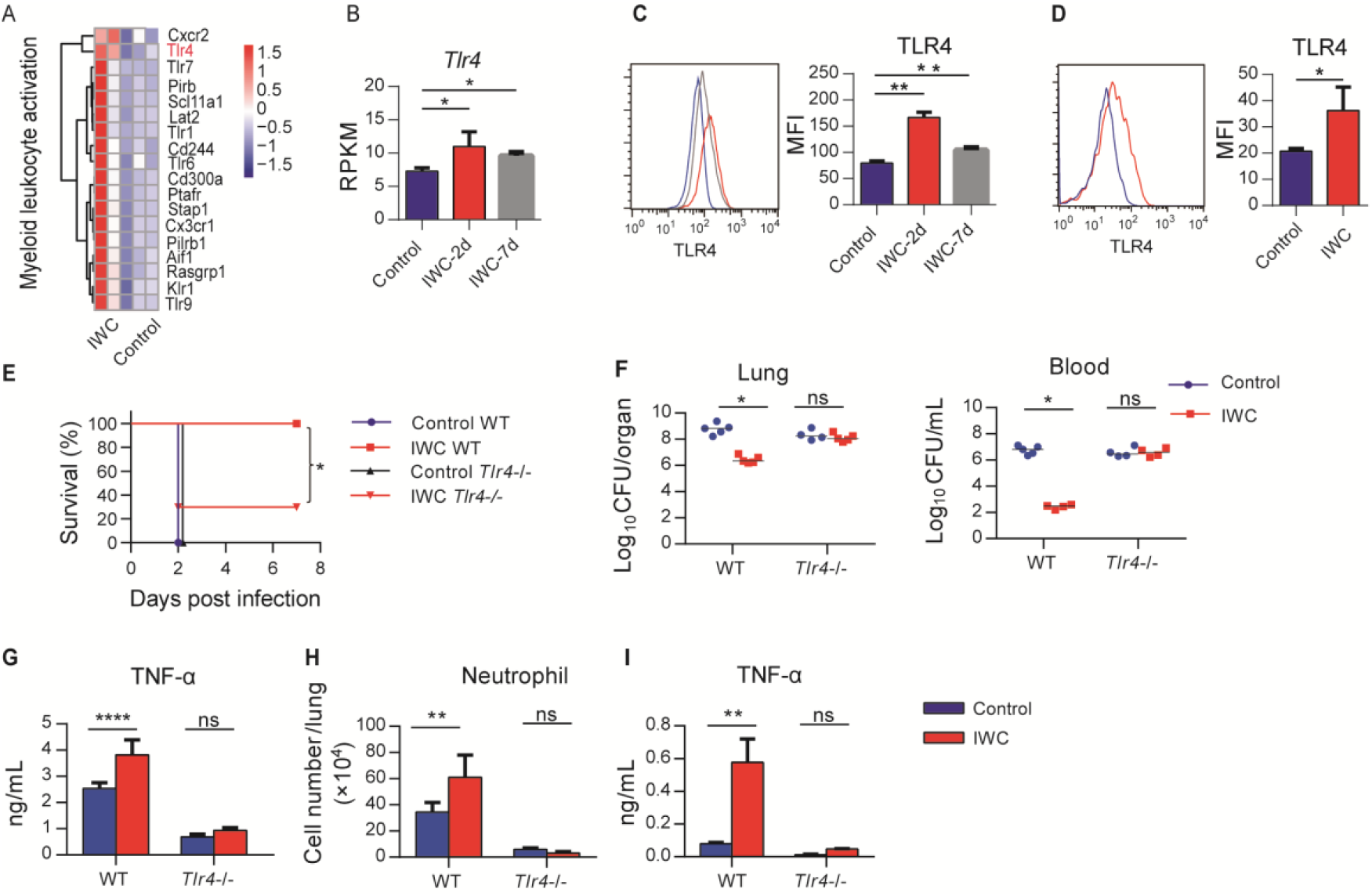
Higher TLR4 expression on IWC-trained AMs mediates rapid protection. (**A**) Heatmap of DEGs associated with myeloid leukocyte activation at day 7 after *A. baumannii* IWC immunization in *Rag1*^-/-^ mice. (**B**) RPKM of TLR4 in lungs on day 2 and day 7 after IWC immunization. (**C**) Representative histogram of TLR4 expression and mean fluorescence index (MFI) of TLR4 on AMs in BALF on day 2 (red line) and day 7 (grey line) after IWC immunization or control (blue line). n=3. For **B** and **C**, \**P* < 0.05, \*\**P* < 0.01, evaluated by ordinary one-way ANOVA. (**D**) Representative histogram of TLR4 expression and MFI of TLR4 on AMs after IWC stimulation *in vitro* (red line) for 7 days. n=4. Data are mean ± SD. \**P* < 0.05, determined by two-tailed unpaired *t* test. (**E-H**) *Tlr4*^-/-^ and WT mice were immunized i.n. with IWC and challenged with *A. baumannii* 7 days later. (n=5-10 mice/ group). (**E**) Survival curve, (**F**) Bacterial burdens at 24 hpi, (**G**) TNF-α in lungs at 2 hpi, and (**H**) Neutrophil infiltration in lungs at 4 hpi. n=4-5 mice. (**I**) TNF-α levels in 2 h culture supernatants of *ex vivo* IWC-stimulated AMs from 7-day vaccinated WT or *Tlr4*^-/-^ mice. For survival, *P* value was calculated by log-rank test. From (F) to (I), \**P* < 0.05, \*\**P* < 0.01, ****, *P*<0.0001, ns, not significant, compared by ordinary two-way ANOVA. In (**F**), the line means median. Data are representative of at least two independent experiments.

### Vaccination-induced rapid protection against *P. aeruginosa* and *K. pneumoniae* infection

Further, we tested whether intranasal vaccination could induce rapid protection in other bacterial pneumonia models. We immunized mice i.n. with IWC of *P. aeruginosa* (IWC(*P*.*a*)), *K. pneumoniae* (IWC (*K*.*p*)), *S. aureus* (IWC(*S*.*a*)) or *S. pneumoniae* (IWC(*S*.*p*)) and challenged with the same bacteria 7 days after immunization. Rapid and efficient protection after intranasal immunization was also observed in *P. aeruginosa* (Figure 6A) and *K. pneumoniae* infected pneumonia models (Figure 6B). However, immunization could not induce effective protection against *S. aureus* (Figure 6C) and *S. pneumoniae* (Figure 6D). Since *A. baumannii, P. aeruginosa*, and *K. pneumoniae* are Gram-negative bacteria and *S. aureus* and *S. pneumoniae* are Gram positive bacteria, we reasoned that intranasal vaccination-induced rapid protection might be an effective way to protect certain Gram negative bacterial pneumonia.

**Figure 6.**
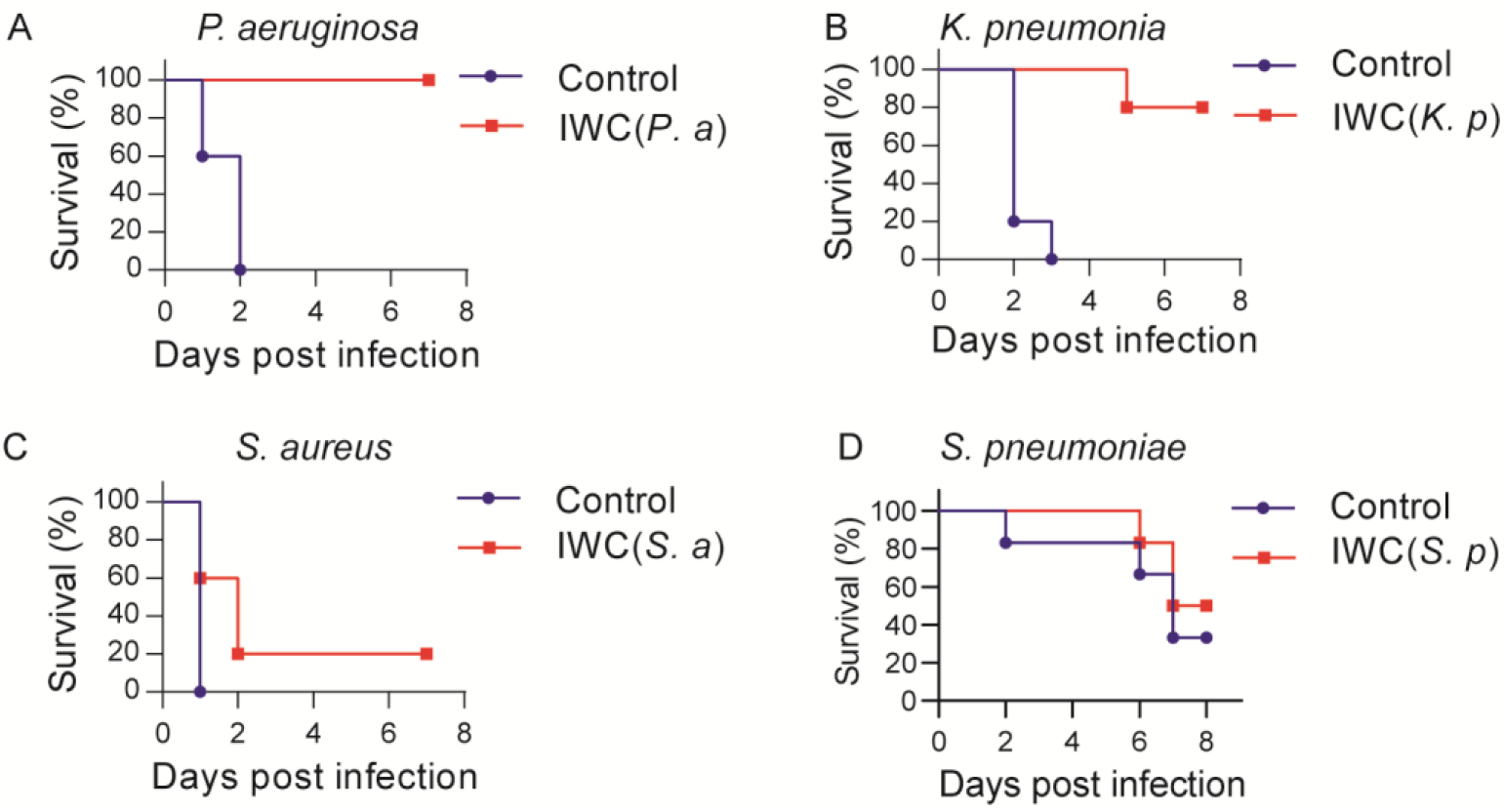
A rapid protection induced by intranasal vaccination against other bacteria. C57BL/6 mice were immunized i.n. with IWC of *P. aeruginosa* (IWC*(P*.*a)*) (**A**), *K. pneumoniae* (IWC*(K*.*p)*) (**B**), *S. aureus* (IWC(*S*.*a*)) (**C**), or *S. pneumoniae* (IWC(*S*.*p*)) (**D**) and were challenged i.t. with the same species of bacteria 7 days later. The survival rates were monitored for 7 days. n=5-10 mice/group. Data are representative of at least two independent experiments.

## Discussion

The challenge of MDR bacterial infection highlights the urgent need to develop rapid-acting vaccine. Currently, only some vector-based virus vaccines are reported to be able to elicit rapid protection by a single dose, such as Ebola and Zika virus vaccines (Marzi et al., 2015; Pardi et al., 2017; Wong, 2019). Here, we showed a single intranasal immunization of IWC of *A. baumannii* elicits a very rapid and complete protection against *A. baumannii* infection 2 or 7 days after vaccination, supported by 100% survival, reduced bacterial burdens, alleviated lung injury, and reduced inflammatory cytokines expression after challenge (Figure 1). What’s more, the vaccination-induced rapid protection is also observed in *P. aeruginosa* and *K. pneumoniae*-infected pneumonia models but not in *S. aureus* and *S. pneumoniae*-infected pneumonia models. Our study suggests the vaccination-induced rapid protection might be a common phenomenon in certain Gram negative bacterial infection, which is critical to protect MDR bacterial pneumonia for inpatients.

The development of immunological memory by vaccination is a central goal in fighting against infections. *A. baumannii* IWC-induced fast protection leads us to suspect whether it is a result of sustained and activated immune response elicited by vaccination. Dynamic response to intranasal immunization of IWC of *A. baumannii* shows that host response rapidly undergoes the priming, resting, and results in memory stage 5 days later (Figure 2A and 3F). The innate immune response 2 days post vaccination might be an activated innate immune response reflected by increase IL-6 and TNF-α response to vaccination (Figure 2A), which provides the host resistance to A. *baumannii* infection. The host response to vaccination completely recovered to baseline level at day 5 post vaccination. However, IWC still induces protection against *A. baumannii* infections at day 7 after immunization, which represents a recall response to vaccination. Upon reexposure to the *A. baumannii* 7 days after vaccination, IWC-vaccinated mice recalls a rapid, heightened TNF-α secretion and chemokine production at 2 h post challenge and subsequently increased neutrophils infiltration earlier at 4 h post challenge in lungs of vaccinated mice (Figure 2B-D). It’s well known that neutrophil, macrophage, and monocytes play essential roles in host defense against *A. baumannii* infection (Qiu et al., 2012; van Faassen et al., 2007). So, vaccination-induced rapid recall responses to infection lead to a rapid elimination of bacteria, thereby limit the uncontrolled inflammation at 24 hpi in vaccinated mice, eventually prevent lung damage and efficiently improve the survival of mice.

For mechanisms study, we highlight the role of trained innate immunity in vaccination-induced rapid protection. Traditionally, immune memory is thought to be an exclusive feature of adaptive immune response of T cells and B cells, which is harnessed to design the vaccine extensively, whereas the role of innate immune response to vaccination is recognized as modulation of adaptive immunity. Recently, trained immunity has been proposed to describe the enhanced immune response of innate cells to second stimuli via epigenetic, metabolic, and functional reprogramming by initial stimulation (Mulder, Ochando, Joosten, Fayad, & Netea, 2019; Netea et al., 2016). Our study in *Rag1*^-/-^ mice (which lack mature T and B cells) further showed innate immune responses can be trained by vaccination to mediate rapid protection (Figure 3). The rapidity of innate response with the trained feature enables it as a good target to design the vaccine to induce rapid protective response against MDR bacteria. AMs are the predominant cells in the airway mucosa and play important roles for infection controls (Hussell & Bell, 2014). The embryonic origin and the ability of self-renewal in steady state of AMs make them be able to store immune memory (Hussell & Bell, 2014). In this study, we found that AMs could be trained by IWC of *A. baumannii* with functional reprogramming, showing increased TNF-α production upon restimulation with the same IWC. More importantly, adoptive transfer of *A. baumannii* IWC-trained AMs into naive recipient mice enhanced the bacteria clearance after lethal challenge with *A. baumannii* (Figure 4), confirming IWC-trained AMs mediate rapid protection. It has been reported that AMs could be trained to gain the memory phenotype in respiratory viral infection models, which is dependent on IFN-γ production from effector CD8^+^ T cells (Yao et al., 2018). However, in our vaccination-trained model, trained AMs is independent on adaptive T cells, since AMs could be trained by IWC directly *in vitro* (Figure 4C) and vaccination-induced protection is still efficient in *Rag1*^-/-^ mice (Figure 3A). It indicates the different mechanisms of AMs training are involved in our models. Although the different mechanisms are involved, our data along with the data of virus-primed AMs both show that AMs could be trained, which can be manipulated to combat MDR bacterial pneumonia and also the respiratory virus infection.

As for how AMs are trained, a range of pattern recognition receptors (PRRs), including Toll like receptors (TLRs), nucleotide-binding oligomerization domain-containing protein 2 (NOD2), and dectin-1 might be engaged to promote trained immunity. Bacille Calmette-Guérin (BCG) and its main component muramyl dipeptide induce trained immunity through NOD2-ligand (Kleinnijenhuis et al., 2012) and β-glucan through dectin-1 receptor (Quintin et al., 2012). Some studies have implicated TLRs are upregulated to BCG or β-glucan training (Kleinnijenhuis et al., 2012; Quintin et al., 2012), however how TLRs are involved in trained immunity is not clear. In this study, we demostrate that TLR4 is elevated on IWC-trained AMs. Enhanced TNF-α production of vaccine-primed AMs upon restimulation is impaired in *Tlr4*^-/-^ mice, which results in reduced protective effect of vaccination (Figure 5). These data suggest that elevated TLR4 expression on AMs might be a trained marker, which can sense the pathogen-associated molecular pattern more efficiently and results in rapid activation of trained AMs compared to naïve AMs. TNF-α is one of the main cytokines which has been thoroughly used as a functional cytokine marker indicating of trained immunity along with IL-6 and IL-1β (Arts et al., 2018). In this study, we found that vaccination-trained immune response produces heightened TNF-α, but not IL-6 and IL-1β when encounter with the infection (Figure 2B). In addition, blocking TNF-α with specific antibody before challenge significantly reduces vaccine-induced rapid protection (Figure 4F). These data collectively indicate that enhanced production of TNF-α from AMs is a functional indicator of trained AMs and responsible for vaccine-elicited rapid protection.

The limitation of this study is that we couldn’t test the vaccination-induced rapid protection in more MDR bacterial pneumonia models, due to the lack of more bacterial pneumonia models in our hands. Also, our study also leaves many open questions, such as how AMs are trained by vaccination, what ligand-receptor pairs are responsible for training AM, what molecular mechanism is involved. Further investigation is needed to better understand the trained immunity of AMs, which in turn will pave the way for improved vaccine design.

In summary, in this study we demonstrate that intranasal immunization of IWC of certain bacteria induces a rapid and sufficient protection against lethal respiratory infection through inducing trained immunity of AMs. Our study highlights the importance and the possibility of harnessing trained immunity of AMs to design rapid effecting vaccine. Even for long-lasting effect of vaccine, exploiting the classical adaptive memory and trained innate immunity in an integrated fashion seems plausible for a potential good design of vaccination strategies against bacterial infection.

## Materials and Methods

### Experimental design and ethical approval

This study was designed to determine whether intranasal immunization of IWC could elicit rapid protection against bacterial pneumonia and explore the underlying mechanisms. For animal studies, 6 to 8 weeks old, female C57BL/6, *Rag1*^-/-^, and *Tlr4*^-/-^mice were used. The animal protocols adhered to the National Institutes of Health Guide for the Care and Use of Laboratory Animals and were approved by the Institutional Animal Care and Treatment Committee of West China Hospital, Sichuan University (Approval No. 2019190A). A MS Excel randomization tool was used to randomize the mice to different treatment groups. To assess the protection efficacy of IWC, survival rate, clinical score, bacterial burdens, and lung histopathology of mice were monitored after infection. RNA-seq, *ex vivo* and *in vitro* AMs stimulation model and adoptive transfer of AMs were performed to investigate the mechanisms underlying the rapid protection. The animal studies were not blinded. The group sizes for survival varied from 5 to 10 in the different studies and 3 to 5 for analysis of immune response. All experiments were conducted at least 2 times independently, which was indicated in figure legends.

### Mice

Female C57BL/6 mice were purchased from Beijing HFK Bioscience Limited Company (Beijing, China). Rag1 gene knockout mice (*Rag1*^-/-^, B6.129S7-Rag1^tm1Mom/J^), TLR4 gene knockout mice (*Tlr4*^-/-^, C57BL/10ScNJNju) and control WT were purchased from Model Animal Research Center of Nanjing University. The mice were kept under specific pathogen-free conditions.

### Bacterial strains

*A. baumannii* strain LAC-4 was kindly provided by Professor Chen (Harris et al., 2013). *P. aeruginosa* strain XN-1, *K. pneumoniae* strain YBQ and *S. pneumoniae* was isolated from Chongqing southwest hospital. *S. aureus* strain KM-22 was isolated from the Second Affiliated Hospital of Kunming Medical University. The bacteria were grown in tryptone soy broth (*A. baumannii*), luria bertani broth (*P. aeruginosa, and K. pneumoniae*), mueller-hinton broth (*S. aureus*), or blood agar plate (*S. pneumoniae)* at 37°C. At mid-log-phase, bacteria were collected and suspended in phosphate buffer saline (PBS). Fresh bacteria were used to infect the mice. For inactivated whole cells (IWC) preparation, fresh bacteria were fixed with 4% paraformaldehyde.

### Intranasal immunization and pneumonia model

Mice were anaesthetized by intraperitoneal injection of pentobarbital sodium (62.5 mg/kg of body weight) and then immunized intranasally (i.n.) with IWC (1×10^8^ CFUs in 20 μl PBS) or PBS as a control. Mice were infected with a lethal dose of bacteria intratracheally (i.t.) through mouth via a soft-end needle under direct visualization to establish pneumonia model (Gu et al., 2018). The survival rate, clinical score, bacteria burdens, and lung pathology were evaluated as described previously (Gu et al., 2018). The lethal doses for different bacteria are as follow: 2×10^7^ CFUs for *A. baumannii*; 1×10^7^ CFUs for *P. aeruginosa*; 5×10^7^ CFUs for *S. aureus*; 2×10^6^ CFUs for *K. pneumoniae; or* 1×10^7^ CFUs for *S. pneumoniae*.

### ELISA

TNF-α, IL-6, and IL-1β concentrations in serum, lung homogenates, and cell culture supernatants were detected using mouse TNF-α ELISA kit, mouse IL-6 ELISA kit, and mouse IL-1β ELISA kit (eBioscience, San Diego, CA, USA) following the manufacturer’s instructions.

### Real-time PCR

Total RNA of lungs was extracted by RNA iso Plus (Takara Biotechnology, Dalian, China) and reverse transcribed to cDNA with PrimeScript™ RT reagent Kit (Takara Biotechnology). Gene expression was detected using SYBR green Premix (Takara Biotechnology) on CFX96 real-time PCR detection machine (Bio-Rad, Hercules, CA, USA) with specific primers listed in Supplementary Table S1. The ΔΔCt method was used to calculate the relative gene expression with β-actin as the housekeep-gene.

### Preparation of BALF and lung cell suspension

Cell were obtained from bronchoalveolar lavage fluid (BALF) as described (Gu et al., 2018). Perfused lungs were cut into small pieces and digested with 1 mg/mL collagenase D (Sigma-Aldrich, St. Louis, MO, USA) and 100 μg/mL DNAase (Sigma) at 37°C for 60 min. Cell suspension was prepared by crushing and filtering the digested tissue through a 70 μm cell strainer (BD Biosciences, New Jersey, USA) and the cell numbers were counted by Countess II Automated Cell Counter (Thermo Fisher Scientific, MA, USA)

### Flow cytometry

Cell suspensions were blocked with rat serum then stained with fluorophore-conjugated specific or isotype control antibodies in the dark at 4 °C for 30 min. The antibodies were as follows: CD45-PE/Cy7 (30-F11), CD11b-PerCP/Cy5.5 (M1/70), F4/80-APC (BM8), Ly-6C-PE (HK1.4), Ly6G-FITC (1A8), CD11c-PE (N418) from Biolegend (San Diego, CA, USA) and TLR4-PE (UT41) from Invitrogen (Carlsbad, CA, USA). Labeled cells were run on a BD FACSCanto™ II flow cytometer (BD Biosciences) and analyzed with FlowJo (BD Biosciences). AMs were defined as CD45^+^CD11b^−^F4/80^+^; monocytes were defined as CD45^+^CD11b^+^ Ly6C^+^; Neutrophils were defined as CD45^+^CD11b^+^Ly6G^hi^. The cell numbers of each cell types were calculated with total cell number multiplied by the cell percentage. AMs in BALF were identified as CD11c^+^ F4/80^+^ cells and TLR4 expression on AMs were detected by flow cytometry and expressed as mean fluorescence index (MFI).

### RNA sequencing (RNA-seq)

Tissue samples from lungs were sent to Wuhan Seqhealth Co., Ltd. (Wuhan, China) for RNA-seq. Briefly, Total RNAs were extracted from lung samples using TRIzol (Invitrogen) and DNA was digested by DNaseI after RNA extraction. A260/A280 was examined with Nanodrop™ One spectrophotometer (Thermo Fisher Scientific) to determine RNA quality. RNA Integrity was confirmed by 1.5% agarose gel electrophoresis. Qualified RNAs were finally quantified by Qubit3.0 with Qubit™ RNA Broad Range Assay kit (Life Technologies, Carlsbad, CA, USA). Total RNAs (2 μg) were used for to prepare sequencing library using KC-Total RNA-seq Library Prep Kit for Illumina® (Wuhan Seqhealth Co., Ltd. Wuhan, China) following the manufacturer’s instruction. PCR products corresponding to 200-500 bps were enriched, quantified and finally sequenced on Hiseq X 10 sequencer (Illumina).

### Analysis of RNA-seq data

Raw data of sequencing were cleaned using Trimmomatic software. The clean reads after quality control were mapped to the mouse genome GRCm38 with STAR software (version 2.5.3a). The reads counts for each gene were calculated using FeatureCounts (version1.5.1) and expressed as RPKM (reads per kilobase per million reads). EdgeR package were used to identify the differentially expressed genes by statistics with an adjusted *P* value < 0.05 and fold change > 1.5. Gene Ontology (GO) and (Kyoto Encyclopedia of Genes and Genomes) KEGG enrichment was done with Kobas (Version 2.1.1). Hierarchical clustering and heatmaps were drawing using pheatmap R package and MA-plot was drawn using EdgeR package.

### *In vitro* AMs training model

BALF cells were cultured in DMEM (containing 10% FBS and 1% penicillin/streptomycin) in plate for 1 h and non-adherent cells were discarded and remaining AMs were stimulated with *A. baumannii* IWC (Multiplicity of infection (MOI) =1) for 24 h, then washout and rest for 6 days. Ams were restimulated with IWC (MOI=1) at day 7 and TNF-α in supernatants at 2 h after restimulation was detected by ELISA. In some experiments, AMs were collected at day 7 to detect TLR4 expression with PE-anti TLR4 antibody (UT41, Invitrogen) by flow cytometry. The expression of TLR4 is showed as MFI.

### Magnetic-activated cell sorting (MACS)

AMs from BALF were sorted by positive selection with an anti-mouse CD11c Microbeads kit (Miltenyi Biotech, Bergisch Gladbach, Germany) according to manufacturer’s instruction. Sorted cells were stained with PE-anti-mouse CD11c antibody (N418) and F4/80-APC (BM8) and analyzed by flow cytometry to check the purity.

### Stimulation of AMs *ex vivo*

Mice were immunized i.n. with IWC or PBS as control. AMs (CD11c^+^) sorted by MACS from BALF at day 7 were stimulated *ex vivo* with IWC (MOI=1) for 2 h and supernatant were collected for measuring TNF-α using ELISA.

### Adoptive transfer of AMs

AMs (CD11c^+^) from BALF of IWC-immunized or control mice at day 7 after immunization were sorted by MACS as described above (purity >95%). Donor AMs (5 × 10^4^) were i.t. transferred into the airways of recipient mice. Recipient mice were challenged with *A. baumannii* LAC-4 (2×10^7^) at 4 h after transfer and bacterial burdens in lungs and clinical score were detected 24 hpi.

### Blocking TNF-α *In vivo*

For neutralizing TNF-α, mice were treated with 200 μg anti-mouse TNF-α (XT3.11 clone, BioXcell, West Lebanon, NH) or rat IgG1 isotype antibody intraperitoneally 1 h before infection, the survival was recorded for 7 days.

### Statistical analyses

Bacterial burdens and clinical score data were expressed as median. Other bar graph data were presented as means ± SD. Survival data were compared by log-rank test. For data more than 2 groups, data were evaluated by ordinary one-way ANOVA followed by Tukey’s multiple comparisons test. Data of two samples with normal distribution were compared by two-tail unpaired *t* test. Mann-Whitney U test was used for comparing data of non-normal distribution (bacterial burdens and clinical score). For grouped data, statistical significance was evaluated by ordinary two-way ANOVA. The software GraphPad Prism version 6.0 was used for all statistical analyses. All comparisons used a two-sided α of 0.05 for significance testing and *P* < 0.05 was considered significant. The specific statistical methods were indicated in the figure legends.

## Data availability

Raw data files for RNAseq have been deposited in the NCBI Gene Expression Omnibus under accession number GEO: GSE141729.

## Acknowledgments

General: We thank Prof. Wangxue Chen for kindly providing the *A. baumannii* strain LAC4 and Dr. Georgina T. Salazar for editing the manuscript.

## Supplemental Information

**Figure 1-figure supplement 1.**
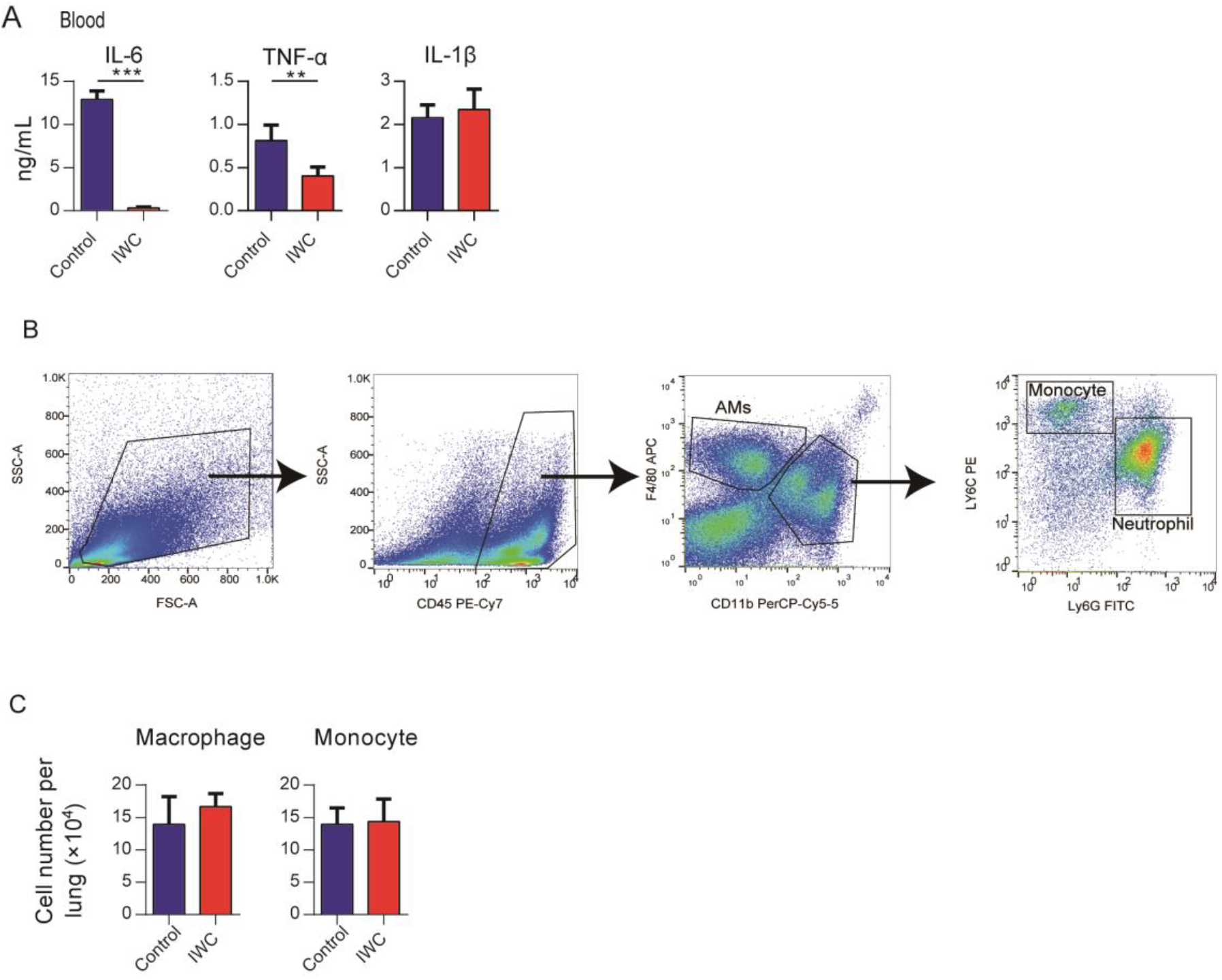
Intranasal IWC vaccination provides rapid protection against *A. baumanii* infection. (**A**) IWC-immunized mice were challenged at day 7 and levels of inflammatory cytokines in blood at 24 hpi were detected by ELISA. (**B**) Gating strategy used in this study for detecting neutrophil, monocyte and alveolar macrophages in lungs. AMs were defined as CD45^+^CD11b^−^F4/80^+^; monocytes were defined as CD45^+^CD11b^+^ Ly6C^+^; Neutrophils were defined as CD45^+^CD11b^+^Ly6G^hi^. (**C**) Numbers of monocytes and macrophages in lungs at 24 hpi were detected by flow cytometry. Data are mean ± SD. *P* value was determined by unpaired *t* test. ** *P* < 0.01, *** *P* < 0.001. Data are representative of two independent experiments (n =4-5 mice/group).

**Figure 3-figure supplement 1.**
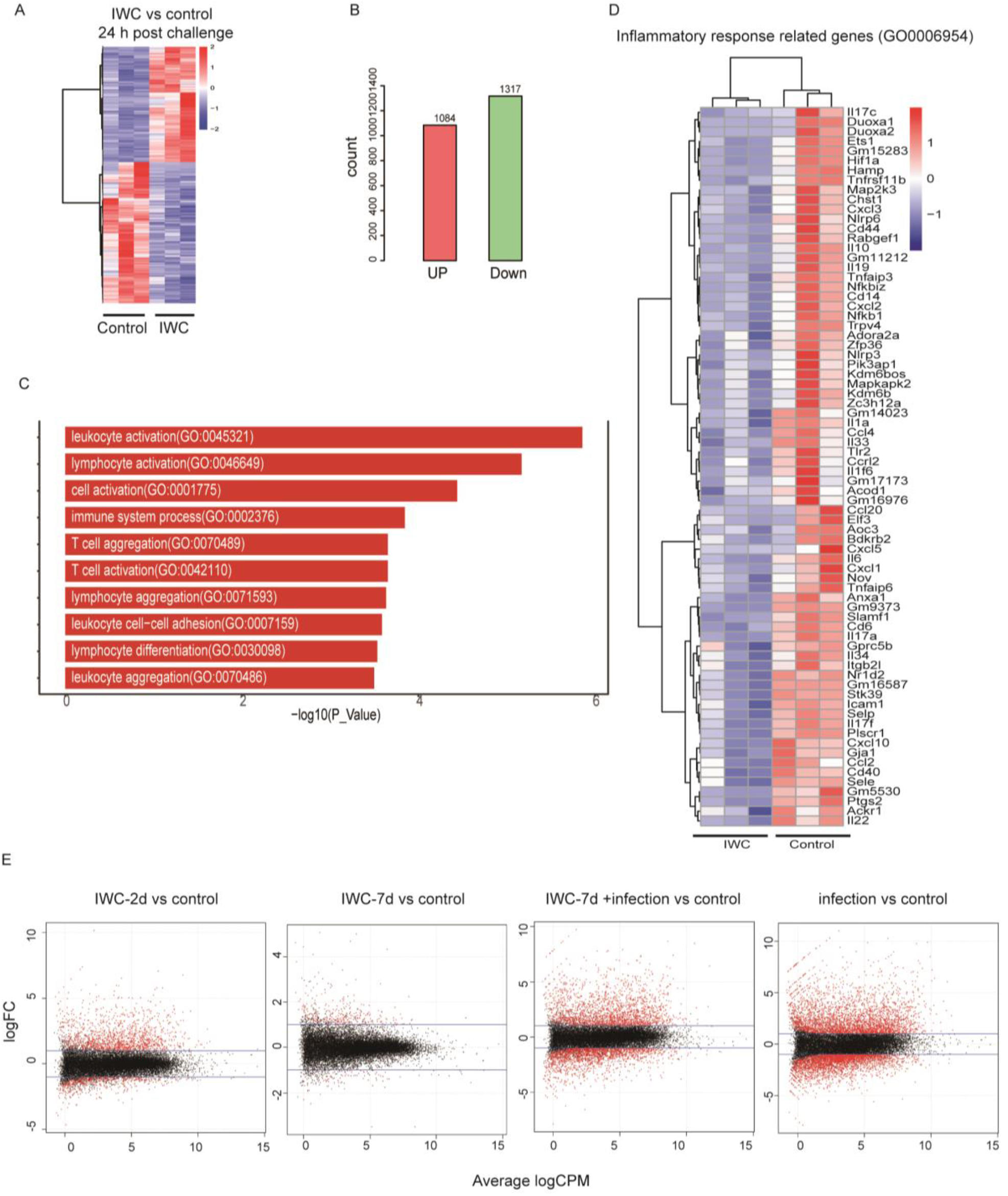
Vaccination-induced protection in *Rag1*^-/-^ mice. (A-D) *Rag1*^-/-^ mice were immunized with IWC of *A. baumannii* or PBS as control and were challenged with *A. baumannii* (2×10^7^ CFU/mice) 7 days later and lungs were processed for RNAseq at 24 hpi. n=3. (**A**) The heatmap of 2401 DEGs in lungs of 7 days immunized and control *Rag1*^-/-^ mice after *A. baumannii* challenge at 24 hpi (n = 3 biological replicates, false discovery rate (FDR) < 0.05). Red indicates increased expression; blue indicates decreased expression. (**B**) Numbers of upregulated and downregulated DEGs of IWC-immunized vs control mice 24 hpi. (**C**) Top 10 GO terms of upregulated DEGs of IWC-immunized group compared to control group at 24 hpi. (**D**) Heatmap of DEGs related to inflammatory response (GO0006954) was shown. False discovery rate (FDR) < 0.05. (**E**) Lung samples from control, IWC-immunized *Rag1*^-/-^ mice at day 2 (IWC-2d), day 7 (IWC-7d), and IWC immunized mice (7 day) at 24 hours after challenge with *A. baumannii* (IWC-7d+infection) and control mice at 24 h after challenge (infection) were processed for RNA-seq. MA plot of DEGs in each treatment group compared with that in control group.

**Figure 4-figure supplement 1.**
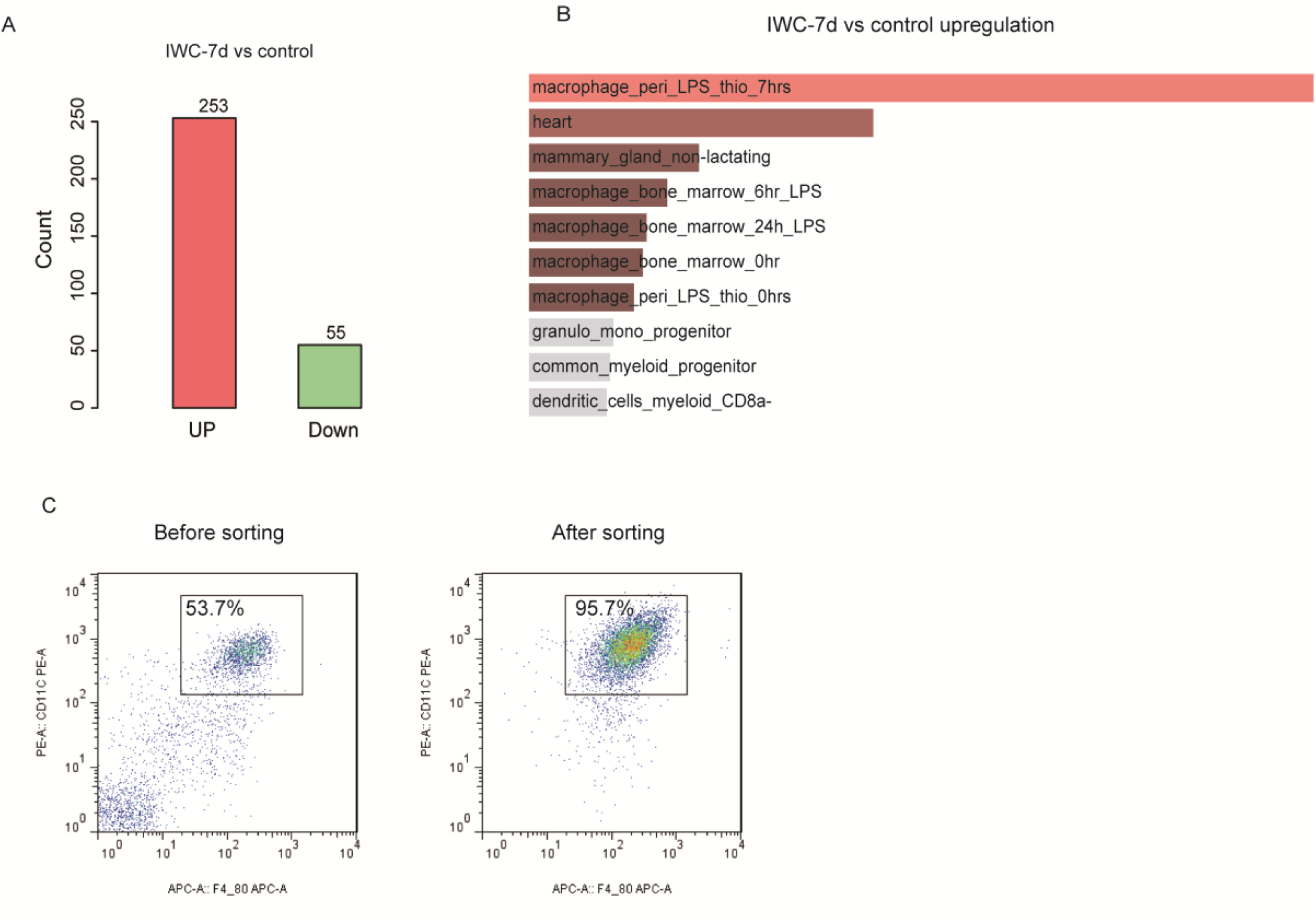
Transcriptional difference at day 7 after immunization. (**A, B**) *Rag1*^-/-^ mice were immunized i.n. with IWC of *A. baumannii* or PBS as control, at day 7 gene expression in lungs from IWC immunized mice (IWC-7d) or control mice were assessed by RNA-seq. (**A**) Numbers of upregulated and downregulated DEGs of lungs in IWC-7d and control mice. (**B**) Top 10 mouse gene atlas terms of upregulated genes in lungs of IWC-7d vs control mice analyzed by Enrichr (https://amp.pharm.mssm.edu/Enrichr/) (*1, 2*). (**C**) Representative flow cytometry of CD11c^+^ cells from vaccinated BALF before and after MACS sorting. CD11c^+^ cells are also F4/80^+^, representing AMs and the purity of AMs after sorting are above 95%.

**Figure 5-figure supplement 1.**
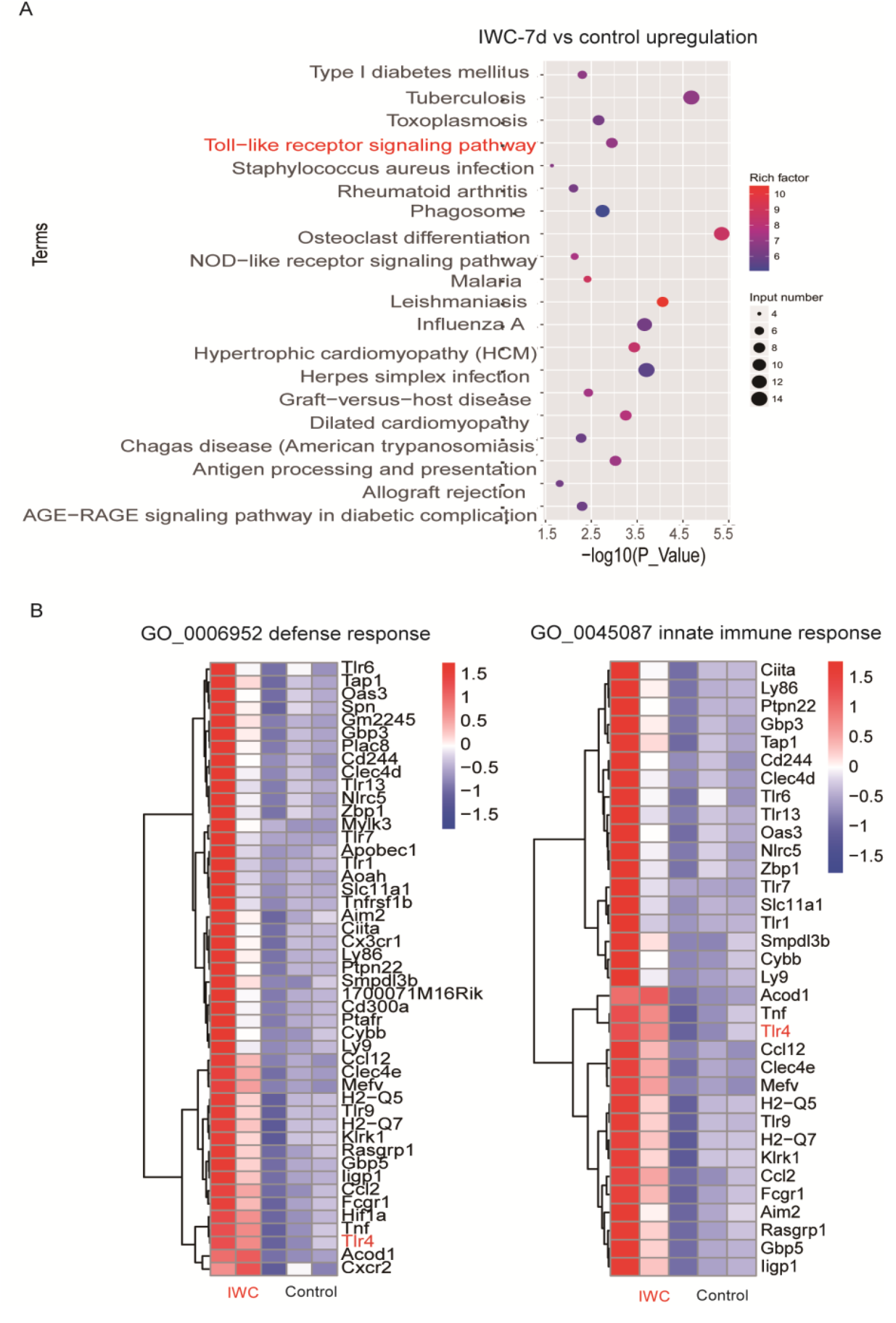
Upregulated differentially expressed genes at day 7 after *A*. IWC immunization in *Rag1*^-/-^ mice. (**A**) Top 20 Kyoto Encyclopedia of Genes and Genomes (KEGG) terms of upregulated DEGs in IWC-7d vs control mice (*P* < 0.05). (**B**) Heatmap of DEGs related to defense response (GO: 0006952) and innate immune response (GO0045087).

**Table S1.**
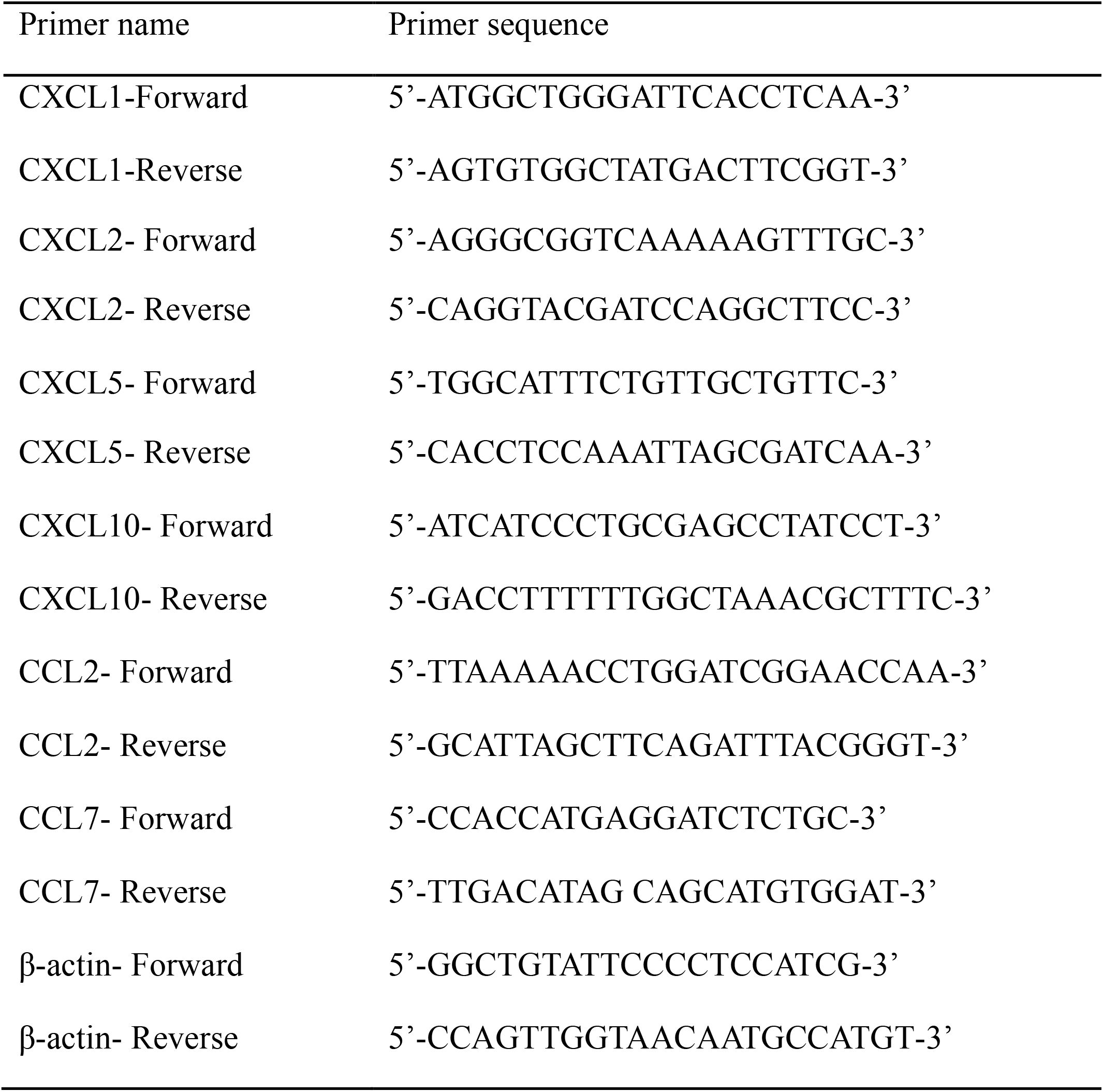
Primers used in real-time PCR.

